# Hypersilencing of SRRM4 suppresses basal microexon inclusion and promotes tumor growth across cancers

**DOI:** 10.1101/2020.03.23.003574

**Authors:** Sarah A. Head, Xavier Hernandez-Alias, Jae-Seong Yang, Violeta Beltran-Sastre, Antonio Torres-Méndez, Manuel Irimia, Martin H. Schaefer, Luis Serrano

## Abstract

RNA splicing is widely dysregulated in cancer, frequently due to altered expression or activity of splicing factors. Microexons are extremely small exons (3-27 nucleotides long) that are highly evolutionarily conserved and play critical roles in promoting neuronal differentiation and development. Inclusion of microexons in mRNA transcripts is mediated by the splicing factor SRRM4, whose expression is largely restricted to neural tissues. However, microexons have been largely overlooked in prior analyses of splicing in cancer, as their small size necessitates specialized computational approaches for their detection. Here we demonstrate that despite having low expression in normal non-neural tissues, SRRM4 is hypersilenced in tumors, resulting in the suppression of basal microexon inclusion. Remarkably, SRRM4 is the most consistently silenced splicing factor across all tumor types analyzed, implying a general advantage of microexon downregulation in cancer independent of its tissue of origin. We show that this silencing is favorable for tumor growth, as decreased SRRM4 expression in tumors is correlated with an increase in mitotic gene expression, and upregulation of SRRM4 in cancer cell lines dose-dependently inhibits proliferation *in vitro* and in a mouse xenograft model. Further, this proliferation inhibition is accompanied by induction of neural-like expression and splicing patterns in cancer cells, suggesting that SRRM4 expression shifts the cell state away from proliferation and towards differentiation. We therefore conclude that SRRM4 acts as a proliferation brake, and tumors gain a selective advantage by cutting off this brake.

**Significance:** Microexons are extremely small exons enriched in the brain that play important roles in neural development. Their inclusion is mediated by the splicing factor SRRM4, also predominantly expressed in the brain. Surprisingly, we find that low expression of SRRM4 outside of the brain is further decreased in tumors, and in fact SRRM4 is the most consistently silenced splicing factor in tumors across tissue types. We demonstrate that SRRM4 inhibits cancer cell proliferation *in vitro* and *in vivo* by inducing a neuronal differentiation program. Our findings add a new element to the overall picture of splicing dysregulation in cancer, reveal an antiproliferative function for SRRM4 and microexons outside of the brain, and may present a common therapeutic intervention point across cancer types.

## Introduction

Alternative splicing (AS) is an important mechanism for increasing the complexity of the human genome, allowing one gene to perform different specialized functions in different cellular or developmental contexts. The most common type of AS in mammalian pre-mRNA is exon skipping, in which a cassette exon is either retained or removed from the mature mRNA transcript. The average exon in humans is ∼140 nucleotides long and contains features that are recognized by splicing factors, which bind both inside and outside the exonic sequence to catalyze the splicing reaction.

Microexons, defined as exons between 3-27 nucleotides in length, have recently been shown to comprise a distinct functional class of cassette exons with higher evolutionary conservation, open reading frame preservation, and enriched localization within protein interaction domains compared with their longer counterparts^1,2^. In contrast to normal exons, microexons are generally too short to contain standard exonic splicing enhancers and thus require a specialized machinery to facilitate their recognition and inclusion. This is mediated primarily by the splicing factor Serine/Arginine Repetitive Matrix 4 (SRRM4, aka nSR100), which has been reported to be expressed at high levels only in neurons^1,3^. SRRM4 plays a critical role in regulating neuronal differentiation, as knockdown and knockout experiments have revealed morphological and functional deficits in cultured neurons as well as the nervous systems of zebrafish and mice^3–7^. In addition, SRRM4 was shown to be downregulated in the brains of some autistic patients, resulting in a global misregulation of microexon splicing^1^, further implicating SRRM4 as a critical factor in brain development. However, the role of microexons in non-neural tissues, if any, has been relatively unexplored to date.

Dysregulation of AS has been implicated in numerous human diseases, including cancer^8^. Loss of splicing fidelity is extremely common in cancer, due either to mutations that directly affect splice sites or regulatory regions within pre-mRNAs, or alterations in splicing factor (SF) expression or activity^9–11^. These changes may result in the expression of protein isoforms that confer selective advantages to cancer cells, either by increasing tumorigenic activity or decreasing tumor suppressive activity. Large-scale studies of AS alterations in cancer have been aided greatly in recent years by consortia such as The Cancer Genome Atlas (TCGA), which has collected multi-omic data from thousands of patient samples. The availability of raw RNA sequencing (RNA-seq) data from both tumor and corresponding normal tissues has facilitated comparative analyses of SF expression and AS, revealing widespread dysregulation of splicing in cancer^12–16^. However, microexons are usually systematically ignored in such analyses, due to their small size and the resulting difficulty in detecting and separating them from background noise. Special computational approaches are therefore required to accurately quantify microexon inclusion^1,17–19^.

Here, we use public data from TCGA to analyze changes in SRRM4 expression and microexon inclusion between tumor and normal samples from 9 different tissues (Figure 1). Surprisingly, we find that not only are SRRM4 and its microexon program globally downregulated in cancer despite having low basal inclusion in normal non-neural tissues, but in fact SRRM4 is the most consistently downregulated across tumor types out of all splicing factors analyzed. We map this decreased expression to a strong increase in methylation of SRRM4 promoters, indicative of epigenetic silencing of SRRM4 expression in cancer. We show that SRRM4 expression and exon inclusion changes anticorrelate with mitotic gene expression in tumors, signifying an inverse relationship with cancer cell proliferation. Correspondingly, we observe marked inhibition of cancer cell proliferation with SRRM4 overexpression both *in vitro* and in a mouse xenograft model, accompanied by induction of neuron-like splicing and expression patterns. We conclude that the microexon splicing program controlled by SRRM4 acts as a brake on proliferation by promoting differentiation, and that tumors gain a proliferative advantage by cutting off this brake.

**Figure 1.**
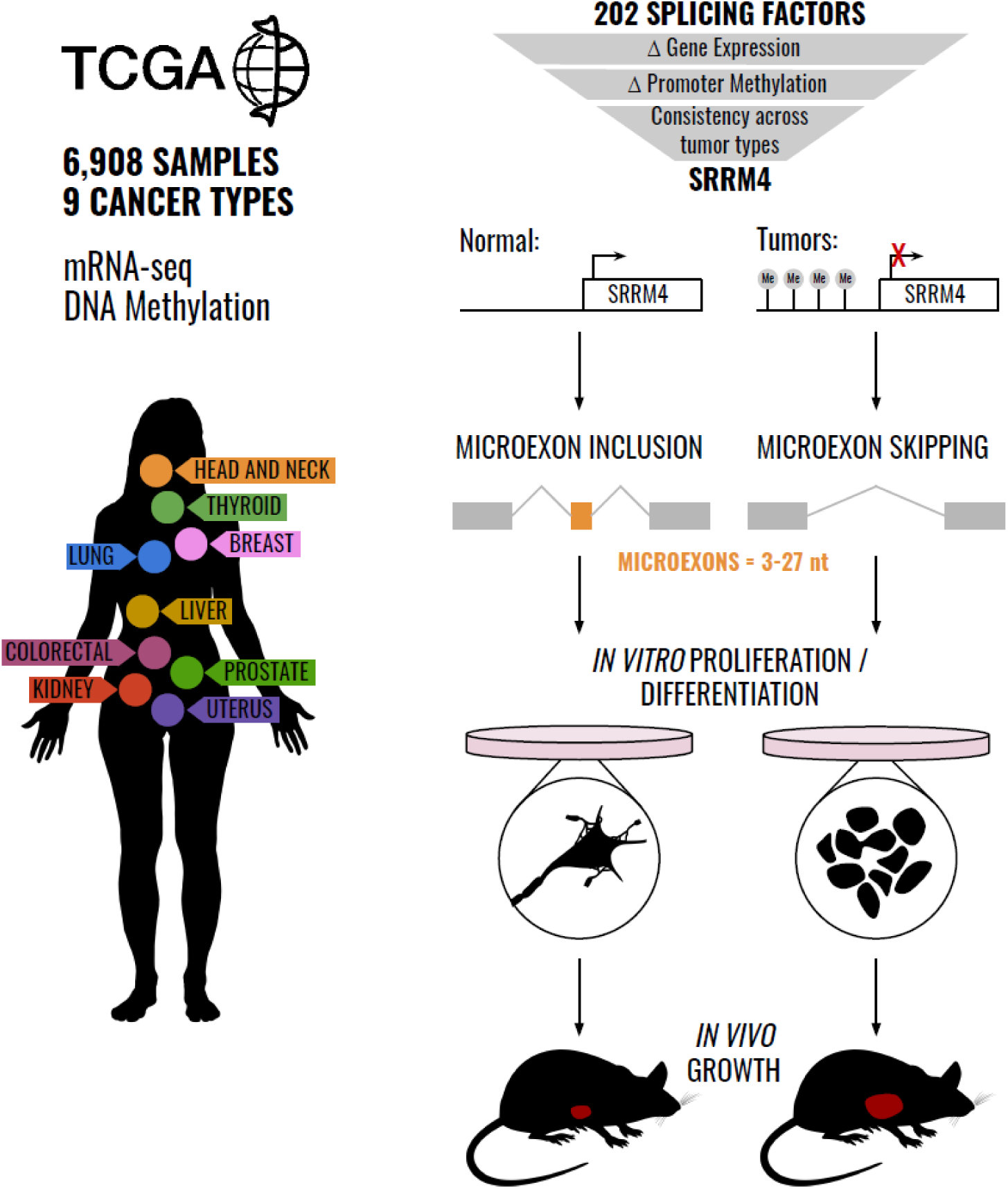
Overview of the study. We analyze splicing factor dysregulation in tumors across 9 different tissues from TCGA. Out of 202 splicing factors, only SRRM4 is consistently silenced by promoter hypermethylation across cancers, resulting in suppressed microexon inclusion in tumors. SRRM4 expression in cancer cells leads to differentiated neuron-like splicing and expression patterns, accompanied by a decrease in cell proliferation *in vitro* and in a mouse xenograft model.

## Results

### SRRM4 expression is silenced in cancer with high consistency across tumor types

Using publicly-available, pre-processed RNA sequencing (RNA-seq) data from TCGA, we first analyzed changes in SRRM4 expression between primary tumor (PT) and solid tissue normal (STN) samples. Nine tissue types were selected that had a sufficiently large number of tumor and matched normal samples to perform conclusive statistics (N≥20 each), as well as available raw RNA-seq files for quantification of splice variants (Table S1). In accordance with the known neural specificity of SRRM4, we found that expression levels in the normal samples were generally low, with median values ranging from 0.0 (liver) to 8.3 (thyroid) RSEM-normalized expression units. Remarkably, the median expression levels of SRRM4 were significantly lower in tumors from all tissues, with the exception of liver that had median expression of 0.0 in both tumor and normal samples (p < 0.001; Wilcoxon-Mann-Whitney test; Figure 2a, Dataset S1). To investigate the mechanism underlying this downregulation, we used DNA promoter methylation array data to assess gene regulation at the epigenetic level^20^, as promoter methylation is a well-established mechanism of gene silencing. Consistent with the downregulated expression, we found an increase in SRRM4 promoter methylation that was highly significant in tumors from all tissues, even in liver (p < 0.001; Wilcoxon-Mann-Whitney test; Figure 2b, Dataset S2). Taken together, these results suggest a global “hypersilencing” of SRRM4 in cancers, beyond its normal silencing in non-neural tissues.

**Figure 2.**
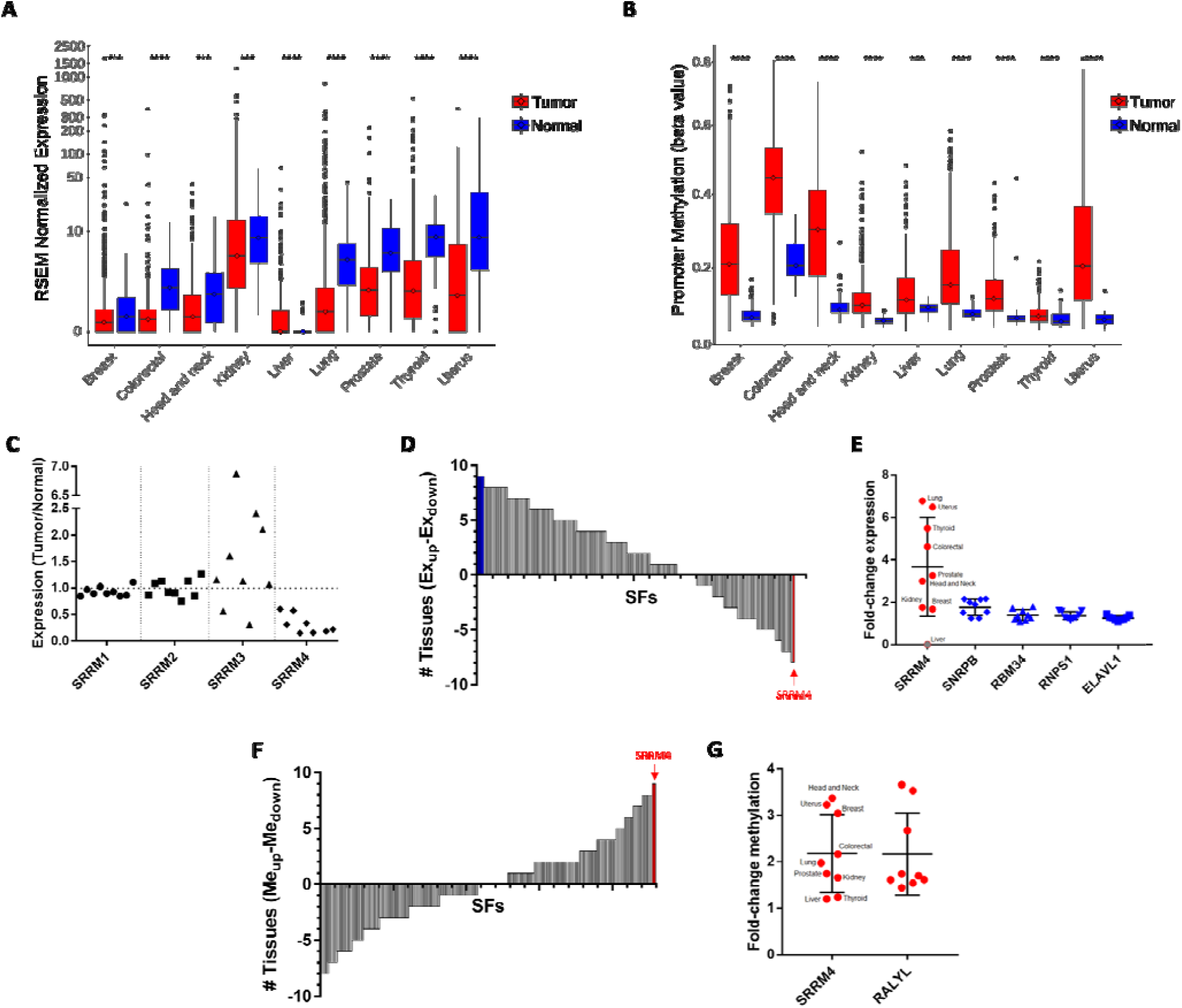
Analysis of TCGA data reveals SRRM4 silencing in cancer across tissue types. A) Changes in SRRM4 expression between normal and tumor samples. *p < 0.05, ** p < 0.01, *** p < 0.001, **** p < 0.0001. B) Changes in SRRM4 promoter methylation between normal and tumor samples. *p < 0.05, ** p < 0.01, *** p < 0.001, **** p < 0.0001. C) SRRM4 is the only SRRM gene family member with consistently changing expression across tissues. Each point in the plot represents the ratio of median expression values between tumor and normal samples for one tissue. D) The sum of the number of tissues with significantly upregulated (positive) and downregulated (negative) expression in tumors, for each of the 202 splicing factors (SFs). E) Fold-change in expression of the four splicing factors that consistently increase across all tumor types (blue) and SRRM4, which decreases in tumors (red), with each point representing one tissue. F) The sum of the number of tissues with significantly upregulated (positive) and downregulated (negative) methylation in tumors, for each of the 202 splicing factors. G) Fold-change in methylation of the two splicing factors that consistently increase across all tumor types (red), with each point representing one tissue.

Despite the high frequency of alternative splicing alterations in cancer, the degree of consistency of SRRM4 silencing across tissues was particularly striking. Other members of the SRRM family (SRRM1-3) did not show the same pattern of consistent change across tissues at either the expression or the promoter methylation level (Figure 2c, Dataset S1-2). In fact, from a list of 202 known splicing factors, only SRRM4 had significantly decreased expression and increased promoter methylation across all tumor types (Figure 2d,f, Dataset S1-2). While four other splicing factors had consistently increased expression (SNRPB, RNPS1, RBM34, ELAVL1), the absolute magnitude of the change in expression was higher for SRRM4 (mean fold-change of 3.67x) compared with the other four (1.25-1.78x) (Figure 2d-e). At the promoter methylation level, one other SF was found to be hypermethylated in all tissues, and to a similar extent as SRRM4 (RALYL; Figure 2f-g), but this hypermethylation did not translate to decreased expression in all tissues. Taken together, these findings demonstrate a global and significant pattern of silencing of SRRM4 in cancer.

### Inclusion of SRRM4 target exons is decreased in tumors

To evaluate the effects of SRRM4 dysregulation at the level of microexon inclusion, we implemented a computational pipeline for the quantification and statistical evaluation of exon inclusion levels from RNA-seq data. Our pipeline makes use of *vast-tools*^21^, a toolset for profiling alternative splicing from sequencing data that is capable of accurately quantifying inclusion levels (defined by “percent spliced in”, or PSI) of exons as small as 3 nucleotides. We ran this pipeline on 6,264 primary tumor (PT) and 644 solid tissue normal samples (STN) from TCGA (Table S1). Inclusion levels were quantified for 219,018 cassette exons, and, for each event, statistical comparisons were performed between tumor and corresponding normal samples in each of the 9 tissues. These results have been uploaded to VastDB^21^ as a public resource (http://vastdb.crg.eu).

For each tissue type, we next defined a list of differentially spliced cassette exons between tumor and normal samples (|ΔPSI| > 5 and q < 0.01, Wilcoxon-Mann-Whitney test; Dataset S3). The resulting list comprised between 468 exons in head and neck cancer to 1,884 in lung cancer, of which between 6.6% (head and neck cancer) and 12.7% (colorectal cancer) were microexons. These differentially spliced exons were significantly enriched for experimentally determined SRRM4 targets (Dataset S4, Methods) in every cancer type tested (p < 10^−6^; Fisher test).

As SRRM4 promotes inclusion of its target exons, we hypothesized that downregulation of SRRM4 in tumors would lead to decreased inclusion levels of known SRRM4-regulated exons^22^. In support of this hypothesis, we found that over 70% of significantly changing SRRM4 target exons across tissues were decreased in tumors compared with normal samples (Figure 3a, Supplementary Figure 1). This result is particularly remarkable given that around two-thirds of known SRRM4-regulated exons are not included in healthy non-neural tissues, meaning they cannot possibly decrease further in tumors (Figure 3b). Of the remaining one-third of SRRM4 target exons with an average PSI > 5 in normal non-neural tissues, a majority (64%) were found to significantly decrease in at least 1 tumor type (Figure 3b, red points). For each individual tissue, the number of SRRM4-regulated exons significantly decreasing in tumors was also larger than those increasing in tumors (Figure 3c). Using all significantly changing exons as a reference, the fraction of decreased SRRM4 target exons was larger than expected by chance in 5 of the 9 tissue types (lung, breast, prostate, uterus, and colorectal; p < 0.05; Fisher test). In agreement with the known role of SRRM4 as a key regulator of microexon splicing, we found that a majority of the differentially included SRRM4 targets were ≤ 27 nucleotides in length. Furthermore, the SRRM4 targets with decreased average inclusion levels in tumors were more highly enriched in microexons (63%) as compared with those increasing in tumors (39%) (Supplementary Figure 2).

**Figure 3.**
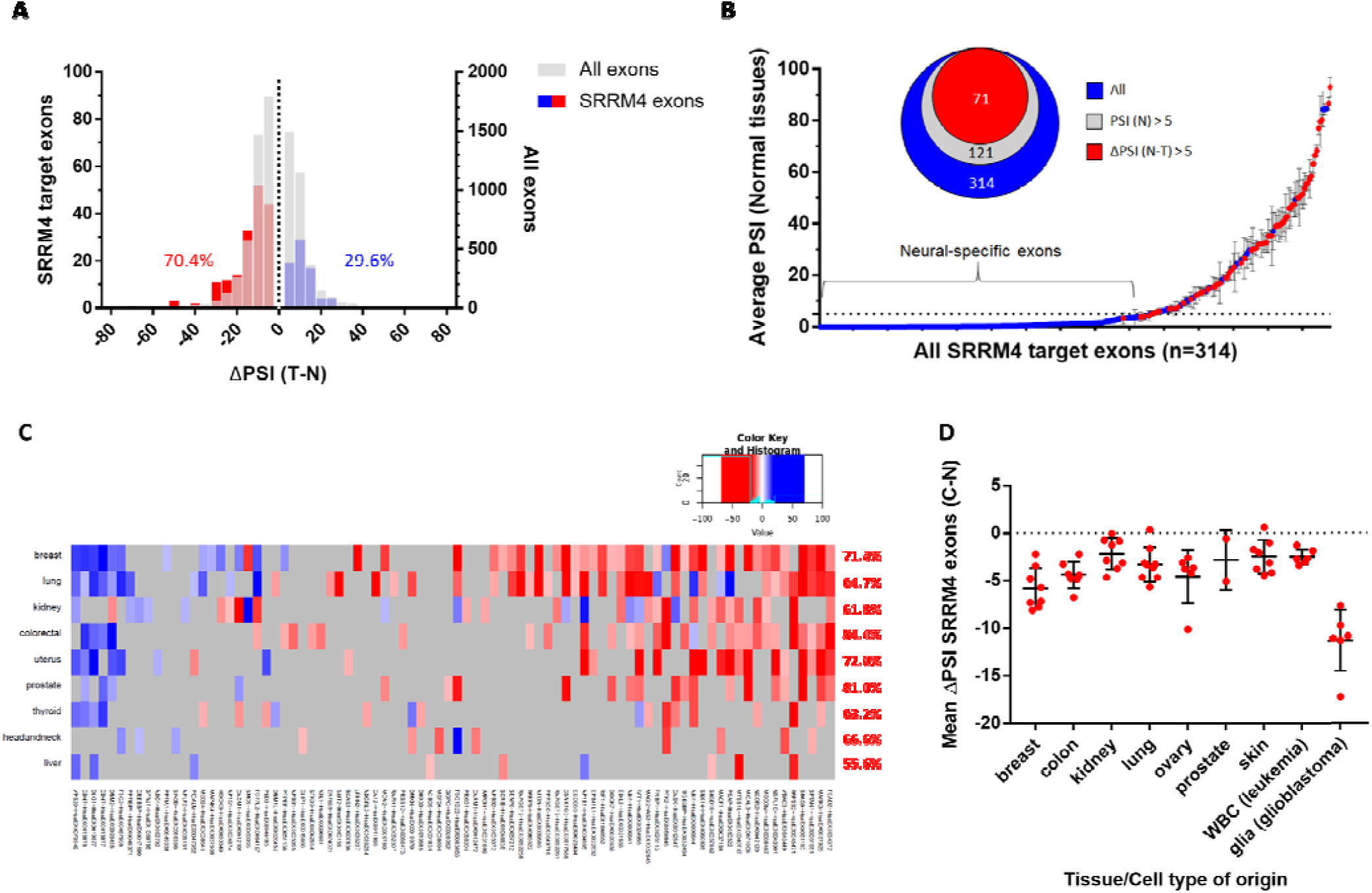
Inclusion of SRRM4-regulated exons is decreased in tumors and cancer cell lines. A) Changes in inclusion levels of SRRM4 target exons (blue/red, left y-axis) compared to non-SRRM4 target exons (grey, right y-axis). B) SRRM4 target exons with respect to their average PSI in normal tissues. Red points are exons with PSI decreasing by at least 5% in at least one tumor type. C) Heatmap of SRRM4-target exons ΔPSIs (Tumor - Normal) across tissue types. Only significantly changing exons (q < 0.01, |ΔPSI| ≥ 5) are shown (red = lower in tumor, blue = higher in tumor). The percentage of SRRM4 target exons decreasing in each tissue type is indicated in red on the right. D) ΔPSI (cell line - normal) of SRRM4-regulated exons in cell lines of the NCI-60 panel. Each point represents the average ΔPSI in one cell line, and cell lines are further grouped by tissue/cell type of origin. Normal data for comparison was from VastDB as indicated (WBC = white blood cells).

To further demonstrate the regulatory impact of SRRM4 activity on SRRM4 target inclusion levels, we correlated the expression levels of SRRM4 with the PSI of its target exons across TCGA tumor samples. The median Spearman correlation was larger than 0 in all cancer types (Supplementary Figure 3a). To determine if the consistently positive correlation levels were larger than expected by chance, we generated background distributions of median correlations between the PSI of SRRM4 target exons and the expression levels of random genes. In 5 of the 9 cancer types we found that the median correlation was larger than expected by chance (lung, breast, prostate, head and neck, and colorectal; p < 0.05; randomization test). Similar results, but in the opposite direction, were obtained when considering the methylation status of SRRM4 instead of its expression (Supplementary Figure 3b). These observations demonstrate that the inclusion of SRRM4 target exons is generally downregulated in tumors as a functional consequence of decreased SRRM4 expression.

To extend the generalizability of our findings, we also monitored the inclusion of SRRM4-target exons in 60 commonly used cancer cell lines from the NCI-60, using a recently published transcriptomic dataset^23^. In agreement with the TCGA results, we found an overall decrease in the inclusion of these exons in cell lines of all tissue/cell types of origin, compared to their corresponding normal samples from VastDB (Figure 3d; Dataset S5). Notably, these cell lines cover 4 cancer types not included in our TCGA analysis: ovarian cancer, melanoma, leukemia, and glioblastoma. In fact, the most striking decrease was observed in glioblastoma, likely due to the higher inclusion of SRRM4 targets in glia than other normal cell types, thereby allowing a greater number of exons the possibility to decrease. Taken together, these results suggest that the observed downregulation of microexon inclusion is a general phenomenon across a wide variety of cancer types.

### SRRM4 expression is inversely related to cell proliferation

To investigate the potential functional consequence of SRRM4 target exon downregulation, we performed GO-term enrichment analysis^24^ on the genes with decreased exon inclusion in tumors. The most significantly overrepresented biological processes among these genes were plasma membrane bounded cell projection organization, generation of neurons, cell morphogenesis involved in neuron differentiation, neuron development, and neuron development (Figure 4a, Dataset S6). These functions are consistent with the known role of SRRM4 in promoting neuronal differentiation, and suggest that decreased expression of SRRM4 might decrease differentiation-related activities in tumors.

**Figure 4.**
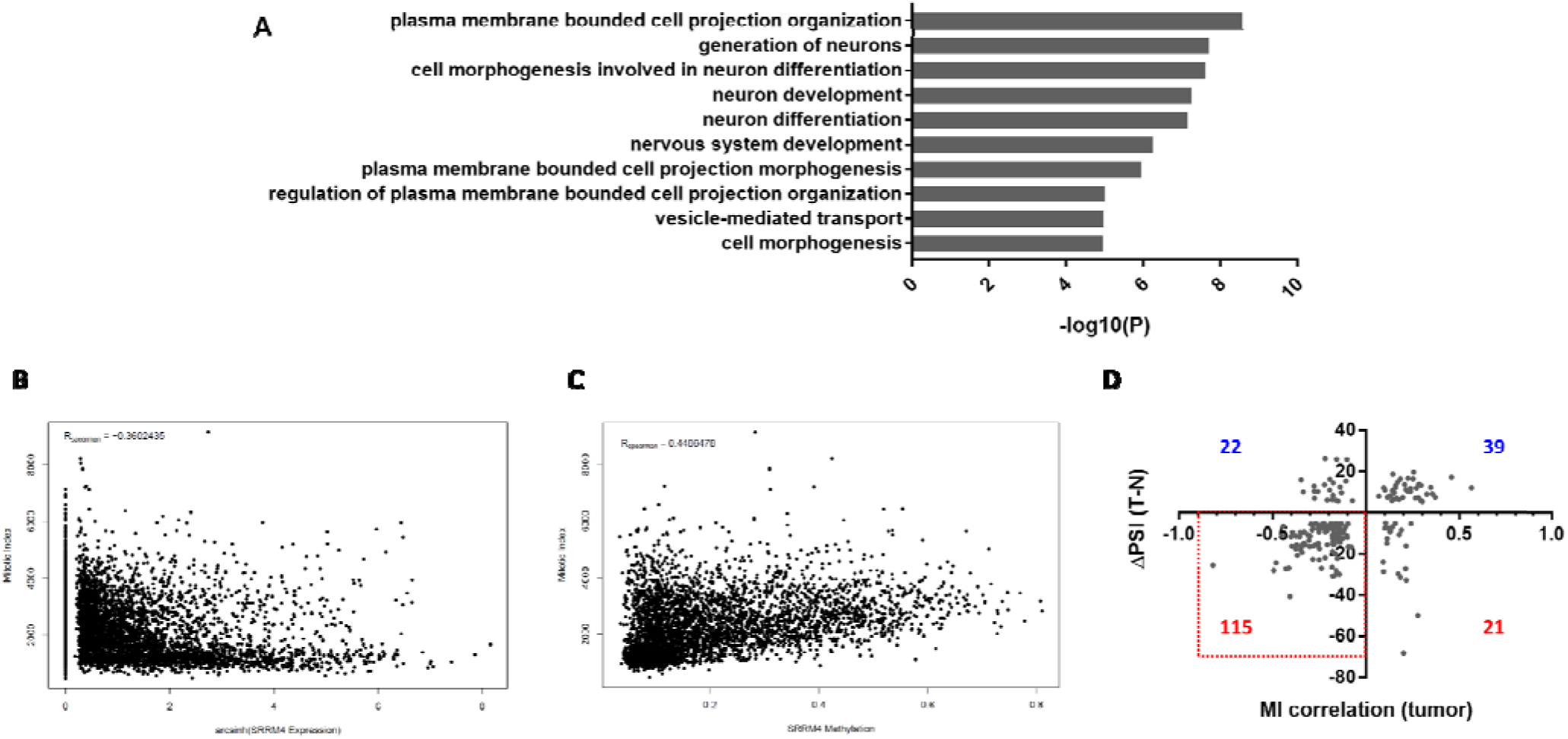
SRRM4 correlates with mitotic index in tumors. A) GO term enrichment of genes with downregulated SRRM4-target exons in tumors from TCGA. The top 10 most significantly enriched terms are shown (GO biological process complete). B) Negative correlation of mitotic index gene signature with SRRM4 expression across TCGA tumor samples. C) Positive correlation of mitotic index gene signature with SRRM4 promoter methylation across TCGA tumor samples. D) Correlation with mitotic index gene signature for each SRRM4 target exon in tumors (x-axis) vs ΔPSI (Tumor-Normal) (y-axis). The number of exons in each quadrant is indicated (red indicates negative ΔPSI; blue indicates positive ΔPSI).

Because neuronal differentiation is accompanied by cell cycle withdrawal and cessation of cell division^25^, we hypothesized that tumors might benefit from downregulating these neuronal differentiation-related processes to gain a proliferative advantage. To assess proliferation in tumor samples, we monitored the mitotic index (MI), an mRNA expression signature that has been shown to correlate with the mitotic activity of cancer cells and is therefore used as an expression-based marker for cell proliferation^26^. In support of our hypothesis, we observed a negative correlation (R_Spearman_ = - 0.36, p < 2.2e-16, Figure 4b) between the MI and SRRM4 expression in tumor samples, while a positive correlation was observed at the promoter methylation level (R_Spearman_ = 0.45, p < 2.2e-16, Figure 4c). This result suggested that SRRM4 expression might negatively affect the proliferation of tumor cells. Furthermore, for each individual SRRM4 exon, we calculated a spearman correlation coefficient between the PSI of the exon and MI across tumor samples, and found a strong negative association between the direction of this correlation for each exon and its ΔPSI in tumors (p = 3.68e-16, binomial test). In agreement, the vast majority (83.9%) of SRRM4 target exons with a negative MI correlation in tumors from a given tissue also had significantly decreased inclusion in tumors from that tissue (Figure 4d), supporting the hypothesis that decreased inclusion of these exons is likely to be associated with increased proliferative activity.

### SRRM4 overexpression inhibits cancer cell growth in vitro and in vivo

To test the hypothesis that SRRM4 expression affects cancer cell proliferation, we generated several stable cell lines with tetracycline-inducible expression of SRRM4. Six different commonly used cancer cell lines were chosen: HeLa (cervical cancer), MCF7 (ER+ breast cancer), MDA-MB-231 (triple negative breast cancer), HCT116 (colon cancer), DU145 (prostate cancer), and SH-SY5Y (neuroblastoma). We also generated control cell lines using empty vector (EV) and an inactive SRRM4 deletion mutant (DM) missing 39 amino acids in the C-terminal enhancer of microexons (eMIC) domain essential for splicing activity^22^. In all six cases, the addition of increasing concentrations of doxycycline to the cells dose-dependently inhibited cell growth only in the WT SRRM4 expressing cells, and not in the EV or DM control lines (Figure 5a-d and Supplementary Figures 5-6). The degree of inhibition was variable between cell lines (Figure 5e), with the greatest antiproliferative effect after 4 days of induction seen in HCT116 (65.8 +/- 4.3% inhibition) and the smallest effect in HeLa (30.2 +/- 8.0% inhibition). Similar anti-proliferative effects were observed in the non-tumor-derived, immortalized kidney cell line, HEK293 (52.7 +/- 6.6% inhibition, 4 days induction) (Supplementary Figures 5-6). Taken together, these results demonstrate that increased SRRM4 expression leads to decreased cell proliferation in both cancer and non-cancer cell lines. These anti-proliferative effects are mediated by SRRM4 splicing factor activity, as evidenced by the fact that the inactive mutant has no such effects.

**Figure 5.**
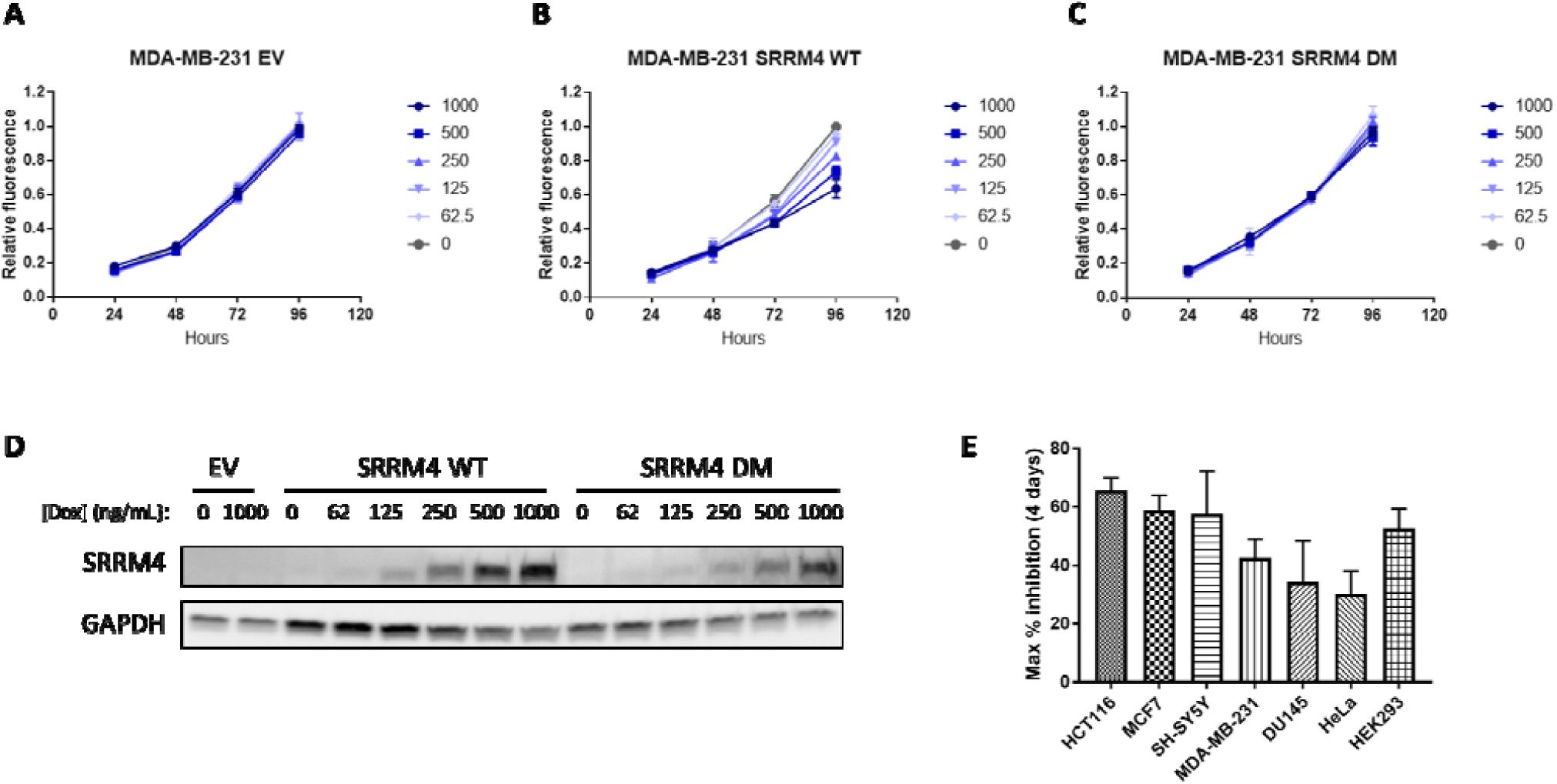
SRRM4 expression inhibits cancer cell proliferation. A-C) Growth curves of MDA-MB-231 transduced with empty vector (EV), wild-type SRRM4 (WT) or deletion mutant SRRM4 (DM), treated with indicated concentrations of doxycycline (ng/mL). Error bars represent standard error of the mean (S.E.M.) of 3 independent experiments. D) Western blot of SRRM4 expression in the same cell lines after 24h induction with doxycycline at the indicated concentrations. Results are representative of 3 independent experiments. E) Maximum % inhibition of cell proliferation after 4 days induction with 1 μg/mL doxycycline in the indicated cell lines transduced with WT SRRM4. Error bars represent standard error of the mean (S.E.M.) of 3 independent experiments.

To further validate the finding that SRRM4 expression affects tumor growth, we performed a mouse xenograft experiment using the MDA-MB-231 cells with inducible SRRM4 expression (WT or DM). Mammary fat pads of athymic female nude mice were implanted orthotopically with the inducible breast cancer cells, and tumors were allowed to grow to 60-80 mm^3^ before induction with 2 mg/mL doxycycline in drinking water (Figure 6a). In agreement with the *in vitro* experiments, induction of WT-SRRM4 tumors with doxycycline significantly decreased the rate of tumor growth compared with uninduced tumors (Figure 6b). In contrast, induction of DM-SRRM4 had no significant effect on tumor growth (Figure 6c). Overall, these results validate the hypothesis that SRRM4 splicing activity can influence tumor growth *in vitro* and *in vivo*.

**Figure 6.**
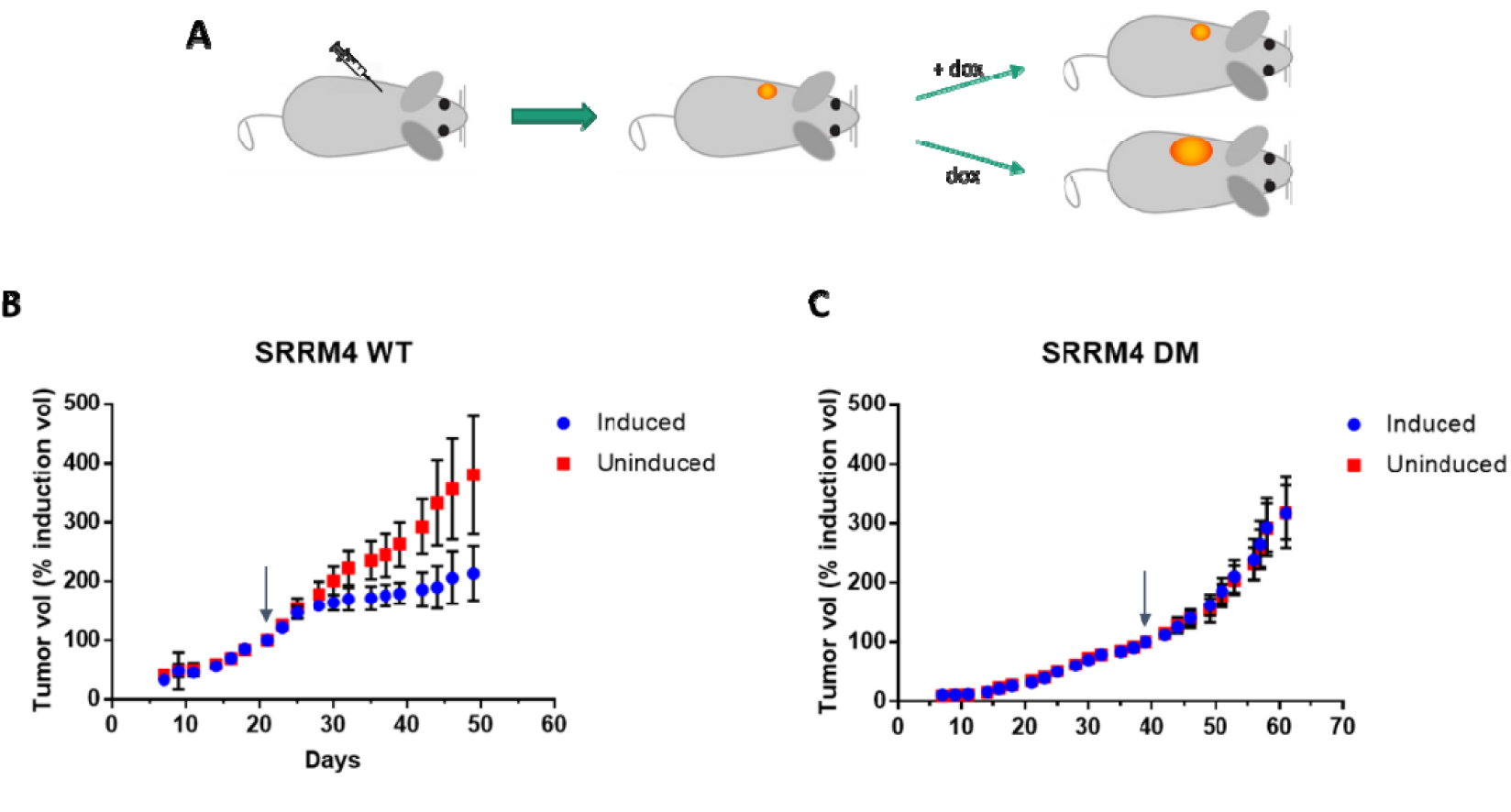
SRRM4 induction inhibits tumor growth in vivo. A) Schematic overview of mouse experiment. Mice were implanted with MDA-MB-231 cells with inducible SRRM4 expression (WT or DM), and tumors were allowed to grow to a certain size before induction with doxycycline in drinking water (2 mg/mL). B) WT SRRM4 tumors grew significantly more slowly in mice receiving doxycycline than non-induced mice. Tumor volume was normalized to tumor size at the time of induction (arrow). Error bars represent standard deviation, n = 6 per group. C) Tumors expressing the inactive SRRM4 mutant (DM) showed no difference in growth rate with or without doxycycline induction. Error bars represent standard deviation, n = 6 per group.

### SRRM4 expression induces neuronal-like expression, splicing patterns, and morphology in cancer cells

To evaluate the functional consequences of altered SRRM4 expression in the above cancer cell lines, we performed RNA-sequencing after 24 h induction of SRRM4 expression. Using the inactive mutant as a reference, we detected increased inclusion (ΔPSI ≥ 25) of between 141 and 216 exons, depending on the cell line (Dataset S7). In total, we found 427 exons with increased inclusion in at least one of the six cell lines, and 69 that were shared between all six cell lines (Figure 7a). Of these, a majority (57%) were shorter than or equal to 27 nt (Figure 7a inset). Although these 69 shared exons generally display low inclusion in normal non-neural tissues compared with neural (Figure 7b), enrichment analysis confirmed that they were significantly overrepresented among the exons decreasing in all 9 cancer types (p < 0.05, one-sided Fisher Exact test), further validating the hypersilencing of SRRM4 target exons in tumors.

**Figure 7.**
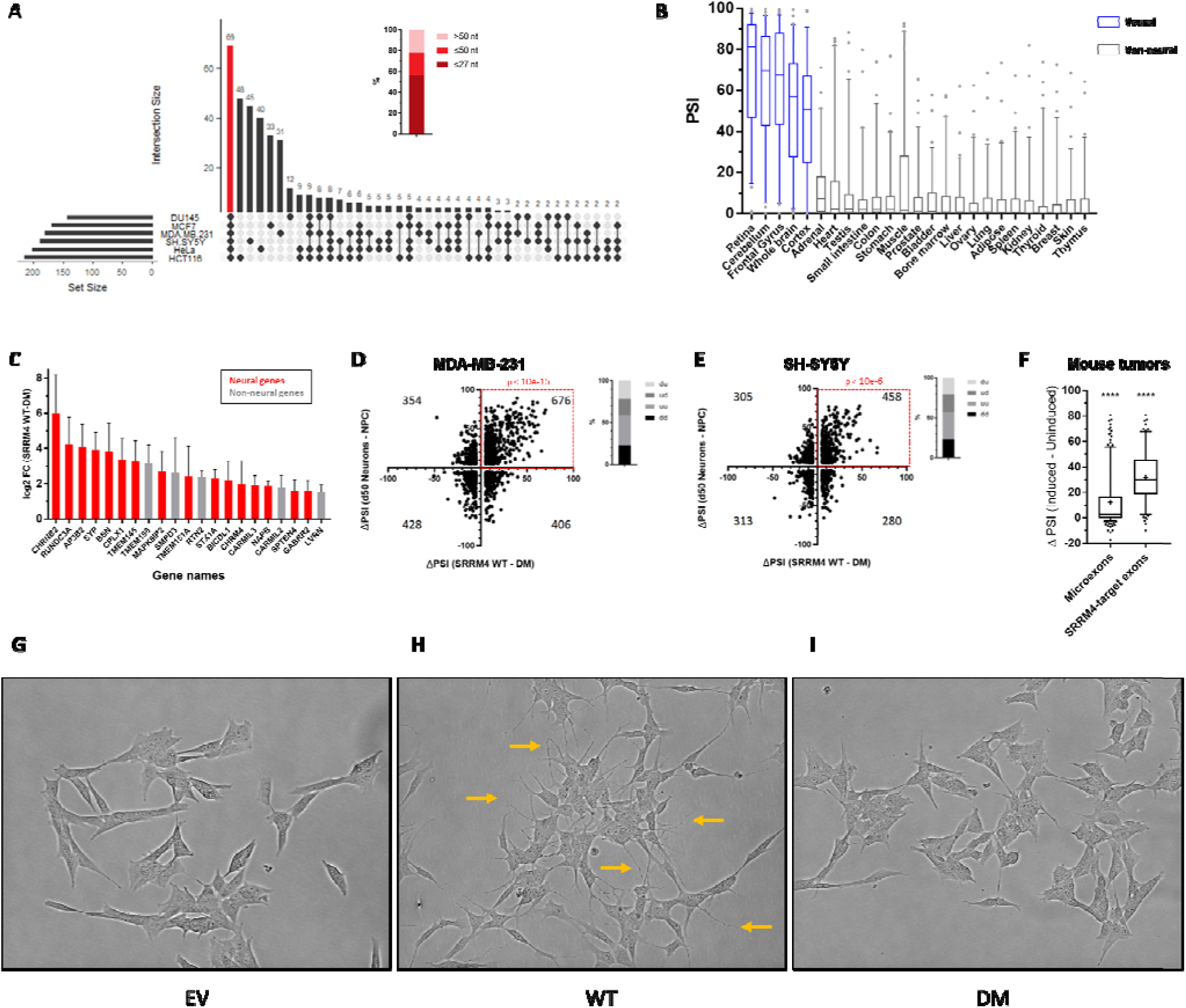
SRRM4 expression leads to neural-like expression and splicing patterns and morphological changes in cancer cells. A) Exons with increased inclusion in all 6 cell lines after SRRM4 induction (ΔPSI ≥ 25). Inset: nucleotide lengths of the 69 shared exons, demonstrating a majority are microexons. B) Box plots of the PSIs of the 69 SRRM4-regulated exons shared between the 6 cell lines across tissue types (from VastDB). C) Genes with increased expression in all 6 cell lines after SRRM4 induction (fold-change ≥ 2). Red bars are genes with known neuronal function (GO: neuron) and/or neural-specific expression patterns. D-E) Overlap between changes in exon inclusion with neuronal differentiation dataset after 7 days of SRRM4 induction in D) MDA-MB-231 or E) SH-SY5Y. F) Changes in inclusion of microexons (≤27 nt) and SRRM4-target exons between the doxycycline-induced and uninduced mouse tumors. The mean ΔPSI for both sets of exons was significantly different than 0 (one-sample t-test, two-tailed; **** p < 0.0001.) G-I) SH-SY5Y cells were transduced with G) empty vector (EV), H) wild-type SRRM4 (WT) or I) deletion mutant SRRM4 (DM). Cell images were taken after 7 days induction with 1 μg/mL doxycycline. WT SRRM4-expressing cells develop numerous projections indicative of neuronal differentiation (orange arrows). Results shown are representative of three independent experiments.

In addition to splicing changes, we also monitored changes in gene expression in the six cell lines after 24 h induction of WT or DM SRRM4 expression, and two of the cell lines (MDA-MB-231 and SH-SY5Y) after 7 days of induction. Compared to the DM control, WT SRRM4-expressing cells showed an upregulation of a large number of neuronal-specific genes. Among the genes found upregulated (≥ 2-fold) in all 6 cell lines after 24 h were those with functions related to neurotransmitter release (AP3B2, BSN, CPLX1, STX1A, SYP), neurotransmitter receptors (CHRM4, CHRNB2, GABRR2) and other genes with neural-specific expression patterns^21^ (CARMIL3, MAPK8IP2, NAPB, TMEM145, TMEM151A, RUNDC3A) (Figure 7c, Dataset S7). We further performed a global comparison of expression changes in the SRRM4-expressing cancer cells (WT vs. DM) with a published dataset of human ESC differentiated into neurons *in vitro* (mature neurons [50 days post-differentiation] vs. proliferating neuronal progenitor cells^27^). This comparison revealed a significant positive association in all cell lines at both 24 h and 7 d induction timepoints (p < 0.005, binomial test; Dataset S8), where a majority of genes with increased expression during neuronal differentiation were also found to increase with WT SRRM4 induction. A similar positive association was also seen for changes in exon inclusion between datasets in all cell lines (p < 10^−5^, binomial test; Figure 7d-e; Dataset S8), in agreement with a previous study comparing splicing differences between neural and non-neural tissues with those seen in the brains of SRRM4 KO mice^5^. The similarity between changes observed in the neuronal differentiation dataset and the cell lines with SRRM4 overexpression further supports the hypothesis that SRRM4 promotes differentiated neuron-like expression and splicing patterns in cancer cells.

We additionally performed RNA sequencing on a small sample of tumors from the mouse xenograft experiment; one uninduced tumor (U19) and two induced tumors (I2 and I14). As expected, both induced tumors had significantly increased inclusion of all microexons, and SRRM4-target exons specifically, compared to the uninduced tumor (Figure 7f, Dataset S9), as well as upregulated expression of neural markers (Dataset S9) similar to what was observed in the cell lines *in vitro* (Dataset S7). Likewise, the differences in both gene expression and exon inclusion between induced and uninduced tumors also showed a highly significant positive association with changes seen in the neuronal differentiation dataset (p < 10^−12^, binomial test; Dataset S8). These results confirm that the differentiation-promoting program induced by SRRM4 has antiproliferative effects on cancer cells *in vivo*.

Consistent with the observed changes in gene expression and exon inclusion, we also observed striking morphological changes in SH-SY5Y upon SRRM4 overexpression, where cells developed numerous dendrite-like projections after 1 week of induction (Figure 7g-i). This morphological change is reminiscent of SH-SY5Y cells that have undergone neuronal differentiation with retinoic acid (RA) and brain-derived neurotrophic factor (BDNF) treatment^28–30^. Indeed, the expression changes observed in both SH-SY5Y and MDA-MB-231 after 7 d induction (Dataset S10) displayed a strong positive association with a published dataset of *in vitro* differentiation of SH-SY5Y cells using RA/BDNF^31^ (p < 10^−7^, binomial test; Dataset S8). While we did not observe such morphological changes in the other cancer cell lines, the consistent upregulation of neural expression and splicing patterns across lines suggests that these cells are shifting towards a neural-differentiation-like state but may not be primed for developing neuronal morphological features, at least after 7 days. However, a recent study reported an increase in cell projections and decreased cell body size in DU145 cells after constitutive long-term SRRM4 overexpression^32^, implying that such changes can occur over longer periods of time.

The above results demonstrate that expression of SRRM4 in cancer cells leads to an upregulation of microexon inclusion, neuron-like expression and splicing patterns, and neuronal morphology, which occurs concomitantly with a decrease in cell proliferation. Taken together, these findings provide a mechanistic explanation for the observed decrease of SRRM4 expression and microexon inclusion in cancer.

## Discussion

In this study, we demonstrate that tumors exhibit downregulated expression of the microexon splicing factor SRRM4 and inclusion of its target exons, with remarkably high consistency across tissues. Our results suggest that this downregulation provides tumors with a growth advantage, as decreased SRRM4 expression in tumors is correlated with an increase in mitotic gene expression, and upregulation of SRRM4 in cancer cell lines dose-dependently decreases proliferation. This antiproliferative activity is also linked to the induction of neuronal differentiation-related genes and splicing patterns, in accordance with the known function of SRRM4 in promoting neuronal differentiation. Proliferation and differentiation are widely considered to be distinct and antagonistic cell states, where terminal differentiation is normally accompanied by exit from the cell cycle and loss of proliferative capacity, and conversely cancer cells evade pro-differentiation programs to promote their proliferation and self-renewing abilities^27,33–35^. We therefore surmise that silencing of SRRM4 provides a selective advantage to cancer cells by shifting the cell state away from differentiation and towards proliferation.

SRRM4 is largely considered to be a neural-specific splicing factor, and as such its basal expression and inclusion of its target microexons are very low outside of the brain. Therefore, our finding that SRRM4 is further hypersilenced in cancers of non-neural origin suggests that low levels of SRRM4 outside of the brain do have a physiologic role as proliferation brakes. Although the effect of downregulating an already lowly expressed gene might be expected to be minimal, any small proliferative advantage would be amplified over time, eventually dominating the cell population. Moreover, despite the fact that around 2/3rds of SRRM4-regulated exons are not included outside of the brain, we observe that a majority of these exons that change in cancer are decreased in tumors, implying that the 1/3rd of SRRM4 exons that are non-zero outside of the brain are sufficient to promote this antiproliferative effect.

While it is well documented that differentiating neurons stop dividing and enter a post-mitotic, terminally differentiated state upon reaching maturity, the molecular mechanisms mediating this proliferation arrest are incompletely understood. Interestingly, mature neurons cannot be induced to reenter the cell cycle without resulting in cell death, and accordingly terminally differentiated neurons are unable to form tumors without first undergoing dedifferentiation^36–40^. It is noteworthy that increased SRRM4 expression and microexon inclusion occur concomitantly with a cessation of cell division during differentiation^5,41,42^. We therefore speculate that SRRM4, in addition to suppressing growth in non-neural tissues, may be involved in mediating proliferation arrest during neuronal differentiation; however, this possibility remains to be explored in future studies.

Unlike alternative splicing events that add or remove entire functional protein domains, microexons are thought of as modulators that “fine-tune” protein activity and protein-protein interaction interfaces^1^. As such, while each individual microexon may have moderate effects on individual processes, modulation of the entire program by the central regulator SRRM4 is a more efficient driver of differentiation by affecting many processes at once. Accordingly, although we find that SRRM4-regulated exons on the whole are downregulated in tumors across all tissues, we do not find any individual SRRM4 exon that changes significantly in every tissue, supporting the idea that the antiproliferative phenotype is likely due to a combined effect of multiple exons that promote neural differentiation. However, we cannot rule out the existence of microexons that did not satisfy our ΔPSI cutoff in all tissues but still have biological significance. Differences across tissues for individual exons could also be due to varying expression levels of the SRRM4 target genes, or additional layers of regulation by other splicing factors. Our ongoing work aims to identify the roles of individual microexons in the antiproliferative and anticancer activity of SRRM4.

Prior studies of SRRM4 in cancer to date have focused exclusively on a rare class of tumors known as neuroendocrine (NE) tumors (0.5-2% of malignancies in adults), due to their shared properties with neural or NE tissues^43^. A series of recent studies have suggested that SRRM4 may play a role in the progression of small-cell lung cancer and NE prostate cancer by inactivating the RE1-silencing transcription factor (REST), in turn promoting expression of neuronal genes^32,44–50^. This program appears to promote transdifferentiation and is associated with poor prognosis in these tumors^44,47,51^. In fact, it has been suggested that SRRM4 mediates increased SOX2 expression, driving prostate cancer cells to a pluripotent phenotype and favoring tumor growth^32^. While we show that SRRM4 overexpression inhibits cell proliferation in all cancer cell lines tested (including the prostate cancer cell line DU145), in our experiments SRRM4 did not induce SOX2 expression (Dataset S7), which could explain the apparent contradiction. NE tumors with SOX2 expression are therefore likely the exception to the here-described selection for silencing of SRRM4 in cancer. Notably, we find that prostate cancers and lung cancers taken as a whole follow the opposite trend as their neuroendocrine subtypes.

Previous studies of alternative splicing alterations in cancer have not specifically taken microexons into account, due to the technical challenges presented by their detection as well as a longstanding underappreciation of their biological relevance. As such, the finding that SRRM4 and its microexon splicing program are silenced in tumors contributes an important new element to the overall picture of splicing dysregulation in cancer. Furthermore, the unusual discovery of such a highly consistent mechanism spanning diverse tissue types may provide novel opportunities for therapeutic intervention in a wide range of cancers. With the advent of splice-switching antisense oligonucleotides that are now being used therapeutically to affect exon inclusion levels^52^, this knowledge could guide the development of new strategies for cancer treatment.

## Materials and Methods

### Statistical Information

Differential expression/methylation of splicing factors and differential exon inclusion were determined by two-tailed unpaired Wilcoxon-Mann-Whitney test. The numbers of samples per group are indicated in Table S1, and p- and q-values are reported in Datasets S1-3. For the distributions shown in Supplementary Figure 3, significance was determined by one-tailed randomization test of the median correlation of SRRM4-regulated events among the corresponding background. For the Gene Ontology analysis, we used the Cytoscape plug-in in ClueGO (v2.5.4). From the list of genes containing downregulated exons, we applied a two-sided hypergeometric test to identify enriched GO terms (between the hierarchical GO level 3 and 5), with Bonferroni step down correction. All terms containing a minimum of 10 genes and constituting at least 1% of the term were included. Binomial tests for associations between changes in different RNA-seq datasets used cutoffs of > 2-fold change in gene expression or ΔPSI > 5 for exon inclusion, and p-values were calculated using the binom.test function in R version 3.4.4. Exact sample sizes and p-values for each pairwise test can be found in Dataset S8.

### TCGA molecular tumor data

We retrieved publicly available and pre-processed RNA-Seq gene expression from firebrowse, and methylation quantification from MethHC^53^. Raw sequencing data in fastq format were retrieved from the GDC legacy archive after obtaining the necessary permissions from dbgap.

### Detection of differentially included microexons in tumors

To determine differential microexon splicing, we analyzed raw RNA-seq files using VAST-TOOLS v2.0, a computational pipeline designed specifically for the quantification of microexon inclusion levels^1^. We ran VAST-TOOLS on 6908 samples (Table S1). Using VAST-TOOLS align we first mapped the reads to the human reference genome hg38 and to quantify PSI levels for each exon and sample. We used VAST-TOOLS tidy to remove exon quantifications with very low quality. We additionally required that PSI levels were determined in at least 20 of both tumor and matched healthy control samples. We then determined differentially spliced microexons by testing for differences in the PSI distributions between tumor and control samples using the Wilcoxon-Mann-Whitney test. We performed FDR correction and considered exons significantly spliced for q-values < 0.01 and an absolute PSI difference between the two distributions of more than 5.

### Correlation between SRRM4 activity and target exon PSI levels

For this study, we generated independent biological replicates for the HEK293 line overexpressing WT SRRM4 as well as a control line overexpressing green fluorescent protein ^22^, and performed RNA-sequencing (Sequence Read Archive accession number PRJNA474911). We used VAST-TOOLS to quantify alternative splicing in these samples and the corresponding matching samples from a previous study ^22^, SRX4193731 and SRX4193721). A list of 314 exons with increased inclusion, i.e. ΔPSI between WT SRRM4 and control greater or equal to 25 in both replicates, was defined as SRRM4 targets and subsequently used in further analyses.

SRRM4 target exon PSI was correlated against the expression and methylation of a set of background genes, in order to determine the significance of SRRM4 correlation by a randomization test. Three different background sets were included: (1) a random set of genes (5000 in expression background, 500 in methylation background), (2) 425 genes that are GO-related to splicing, and (3) 233 genes that are reported to regulate microexons ^54^.

### Plasmids

The cloning of human WT and deletion mutant (ΔPFAM) SRRM4 was described previously ^22^. SRRM4 was amplified by PCR using attB-containing primers (F: 5’-GGGGACAAGTTTGTACAAAAAAGCAGGCTTCATGGCGAGCGTTCAGCAAGGC-3’; R: 5’-GGGGACCACTTTGTACAAGAAAGCTGGGTCTTAGCGCCTCGTGCTGGAGTAG-3’) and reacted with the gateway donor vector pDONR223 by using BP clonase II (11789020, ThermoFisher Scientific, Madrid, Spain). The resulting entry clones containing SRRM4 WT and DM were reacted with the gateway destination vector pCW57.1 (a gift from David Root, addgene plasmid #41393) using LR clonase II (11791020, ThermoFisher Scientific, Madrid, Spain).

### Cell lines and lentivirus production

HEK293 cells were purchased from ATCC (CRL-1573). DU145 were a gift from Luciano Di Croce (CRG). All other cell lines were obtained from the CRG cell line repository. The sex of each cell line is as follows: DU145 Male; HeLa Female; HEK293 Female; MCF7 Female; MDA-MB-231 Female; SH-SY5Y Female; HCT116 Male. All cell lines were cultured at 37°C and 5% CO2 in DMEM (H3BE12-604F/U1, Lonza Group AG [Cultek S.L.U, Madrid, Spain]) supplemented with 10% tetracycline-free FBS (631106, Clontech, [Conda Laboratories, Torrejon de Ardoz, Spain]) and 1% penicillin/streptomycin (15140122, Life Technologies S.A., [Fisher Scientific, Madrid, Spain]). To generate lentivirus, low passage HEK293T cells were plated in 6-well plates at 800,000 cells/well in 3 mL media. The following day, cells were co-transfected with 0.9 μg pCW57.1 (empty vector, SRRM4 WT, or SRRM4 DM), 0.6 μg psPAX2 (a gift from Didier Trono, addgene plasmid #12260) and 0.3 μg pMD2.G (a gift from Didier Trono, addgene plasmid #12259) per well using Lipofectamine 2000 transfection reagent (11668027, ThermoFisher Scientific, Madrid, Spain) according to the manufacturer’s instructions. After 48-72 h, viral supernatant was harvested, filtered through a 0.45 um PVDF filter (SLHV033RS, Merck Chemicals and Life Science S.A, Madrid, Spain), and transferred to the target cell line growing in a 6-well plate at ∼70% confluence. After 48 h, the target cells were trypsinized and replated in 6-cm dishes in media containing 2 μg/mL Puromycin (P8833, Sigma-Aldrich Quimica S.A., Madrid, Spain). Selection media was changed every 48-72 h for a minimum of 1 week before cell lines were frozen in 50% FBS and 10% DMSO and stored in liquid nitrogen for later use.

The generation of HEK293 with inducible WT SRRM4 expression is described elsewhere ^22^.

### Resazurin growth assays

Resazurin sodium salt (R7017, Sigma-Aldrich Quimica S.A., Madrid, Spain) stock was prepared at 0.2 mg/mL in PBS and sterilized by filtration through a 0.22 um PVDF filter (SLGP033RS, Merck Chemicals and Life Science S.A, Madrid, Spain). Cells were seeded in 4 flat-bottom 96-well plates (353072, Corning B.V. [Fisher Scientific, Madrid, Spain]) at a density of 1,000 cells/well in 200 μL of tetracycline-free media, containing 2-fold serial dilutions of doxycycline (D9891-1G, Sigma-Aldrich Quimica S.A., Madrid, Spain) (1000-62.5 ng/mL final concentration) with each condition in triplicate. Every 24 hours, resazurin was added to one plate (10 μL/well) and fluorescence intensity was measured after 5 h using an Infinite M-Plex plate reader from Tecan (Ibérica Instrumentación S.L, Barcelona, Spain) (ex = 560 nm, em = 590 nm). Within each experiment, values were normalized to the mean fluorescence of uninduced cells at the 96 h timepoint. Three replicate experiments were performed for each cell line. The results were graphed using GraphPad Prism v. 7.04 and are presented as normalized means with error bars representing the standard error of the means.

### Western blotting

Cell extracts were harvested by direct addition of 2x SDS sample buffer (1610747, BioRad Laboratories, Alcobendas, Spain) containing 100 mM DTT to the culture plate and incubation on ice for 10 minutes, followed by collection in 1.5 mL eppendorf tubes and denaturation for 5 min at 95°C. Lysates were subjected to SDS-PAGE in Criterion™ TGX Stain-Free™ Precast Gels (567-8124, BioRad Laboratories. Alcobendas, Spain) and transferred to nitrocellulose membranes (IB301001, ThermoFisher Scientific, Madrid, Spain) by iBlot Dry Blotting System (ThermoFisher Scientific, Madrid, Spain). Membranes were blocked in 5% milk in TBS with 0.05% Tween-20 for 1 h at RT before addition of primary antibodies (SRRM4, HPA052783, Sigma-Aldrich Quimica S.A., Madrid, Spain; GAPDH, sc-47724, Santa Cruz Biotechnology) diluted 1:1,000 in blocking buffer, followed by overnight incubation at 4°C. Membranes were washed 3 × 5 min in TBS-T, followed by 1 h incubation at RT in secondary antibodies (Anti-Rabbit: A0545; Anti-Mouse: A6782, Sigma-Aldrich Quimica S.A., Madrid, Spain) diluted 1:10,000 in blocking buffer, and 3 more 5 min washes with TBS-T. Finally, membranes were developed by the addition of HRP substrate (34096, Thermo Scientific, [Fisher Scientific, Madrid, Spain]) and exposed using a LAS-3000 Imaging System from Fuji.

### RNA extraction and sequencing

Cell extracts were harvested by direct addition of QIAzol lysis reagent (79306, Qiagen [WERFEN ESPAÑA, S.A.U. L’Hospitalet de Llobregat, Barcelona]) to the culture plate and incubation for 5 min at RT followed by collection in 1.5 mL eppendorf tubes. RNA was isolated using the miRNeasy mini kit (217004, Qiagen [WERFEN ESPAÑA, S.A.U. L’Hospitalet de Llobregat, Barcelona])) according to the manufacturer’s instructions. For isolation of RNA from mouse tumor tissue, tumors frozen in RNAlater stabilization solution (AM7020, ThermoFisher Scientific, Madrid, Spain) were thawed in QIAzol and disrupted using a Polytron PT1200E homogenizer (KINEMATICA INC, VWR), before continuing with the miRNeasy protocol. Purified RNA was stored at −80°C before sequencing.

RNA quality control, library preparation and sequencing were performed by the CRG Genomics Unit. RNA concentration and quality were assessed by nanodrop and bioanalyzer. An average of 90 million 125-nucleotide paired-end reads were generated for each sample. Raw sequencing data were submitted to the Sequence Read Archive (accession number PRJNA551123).

### Mouse xenograft model

Mouse experiments were performed by Axis Bioservices (Coleraine, Northern Ireland). All protocols were approved by the Axis Bioservices Animal Welfare and Ethical Review Board, and all procedures carried out under the guidelines of the Animal (Scientific Procedures) Act 1986. Female athymic nude mice aged 5-7 weeks weighing approximately 23-30 g were bred in house and housed in IVC cages (up to 5 mice per cage) with individual mice identified by tail mark. Holding conditions were maintained as follows: room temperature 20-24°C, humidity 30-70%, and 12 h light/dark cycle. All animals had free access to standard a certified commercial diet and water, and cages were changed once a week with food and water replaced when necessary. Animals were randomly allocated to treatment groups using Graphpad. MDA-MB-231 cells with inducible SRRM4 expression, generated as described above, were cultured in DMEM with 10% tetracycline-free FBS, 1% penicillin/streptomycin, and 1 μg/mL puromycin. Cells were orthotopically implanted into the inguinal mammary fat pad of the mice using a 25G needle (5×10^6^ cells in Matrigel 1:1, one tumor implanted per mouse). When tumors reached approximately 60-80 mm^3^, mice were randomly assigned to treatment groups using Graphpad, and treated with or without doxycycline diluted to a concentration of 2 mg/mL in drinking water. Tumor volume was measured 3 times per week using digital calipers, and measurements continued for 4 weeks after induction with doxycycline. The length and width of the tumor was measured and volume calculated using the following formula: volume = (length x width^2^)/2. The bodyweight of all mice was measured and recorded 3 times per week, and mice were observed daily for any signs of distress or changes to general condition. After 28 days post doxycycline induction, animals were euthanized via carbon dioxide inhalation, and tumors were resected and weighed before being placed into RNAlater stabilization solution and frozen at −80°C for later analysis.

## Acknowledgements

We acknowledge the support of the Spanish Ministry of Economy and Competitiveness, ‘Centro de Excelencia Severo Ochoa’, the CERCA Programme / Generalitat de Catalunya, and the Spanish Ministry of Economy, Industry and Competitiveness (MEIC) to the EMBL partnership. S.A.H. is supported by a Marie Skłodowska-Curie Individual Fellowship from the European Union’s Horizon 2020 research and innovation programme (MSCA-IF-2017-794629). X.H. is supported by a PhD fellowship from the Fundación Ramón Areces. The results published here are in part based upon data generated by the TCGA Research Network: https://www.cancer.gov/tcga. We thank the CRG Genomics Unit for assistance with RNA sequencing services. We thank Patryk Polinksi, Reza Sodaei and Sophie Bonnal for advice and provision of reagents, Juan Valcarcel for critical reading of the manuscript, and Joseph DiBartolo for assistance with data processing.

## Author Contributions

Conceptualization, M.H.S. and L.S.; Methodology, S.A.H, M.H.S. and L.S.; Investigation, S.A.H., V.B-S., and A.T-M.; Software, J.S.Y., X.H., M.H.S. and M.I.; Formal Analysis, S.A.H., X.H. and M.H.S.; Writing – Original Draft, S.A.H. M.H.S. and L.S.; Writing – Review and Editing, S.A.H., M.I., M.H.S., L.S.; Funding Acquisition, S.A.H. and L.S.; Supervision, M.H.S. and L.S.

## Materials and correspondence

Further information and requests for reagents may be directed to Luis Serrano.

## Data availability

Raw RNA sequencing data generated in this study have been deposited in the NCBI Sequence Read Archive (SRA) with accession numbers PRJNA474911 and PRJNA551123. Processed RNA sequencing data, including gene expression and exon inclusion quantification, are included in this published article as supplementary datasets.

