## Supplemental information for "Hypersilencing of SRRM4 suppresses basal microexon inclusion and promotes tumor growth across cancers"

**This PDF file includes:**

Figures S1 to S5

Table S1

Legends for Datasets S1 to S10

SI References

**Other supplementary materials for this manuscript include the following:**

Datasets S1 to S10

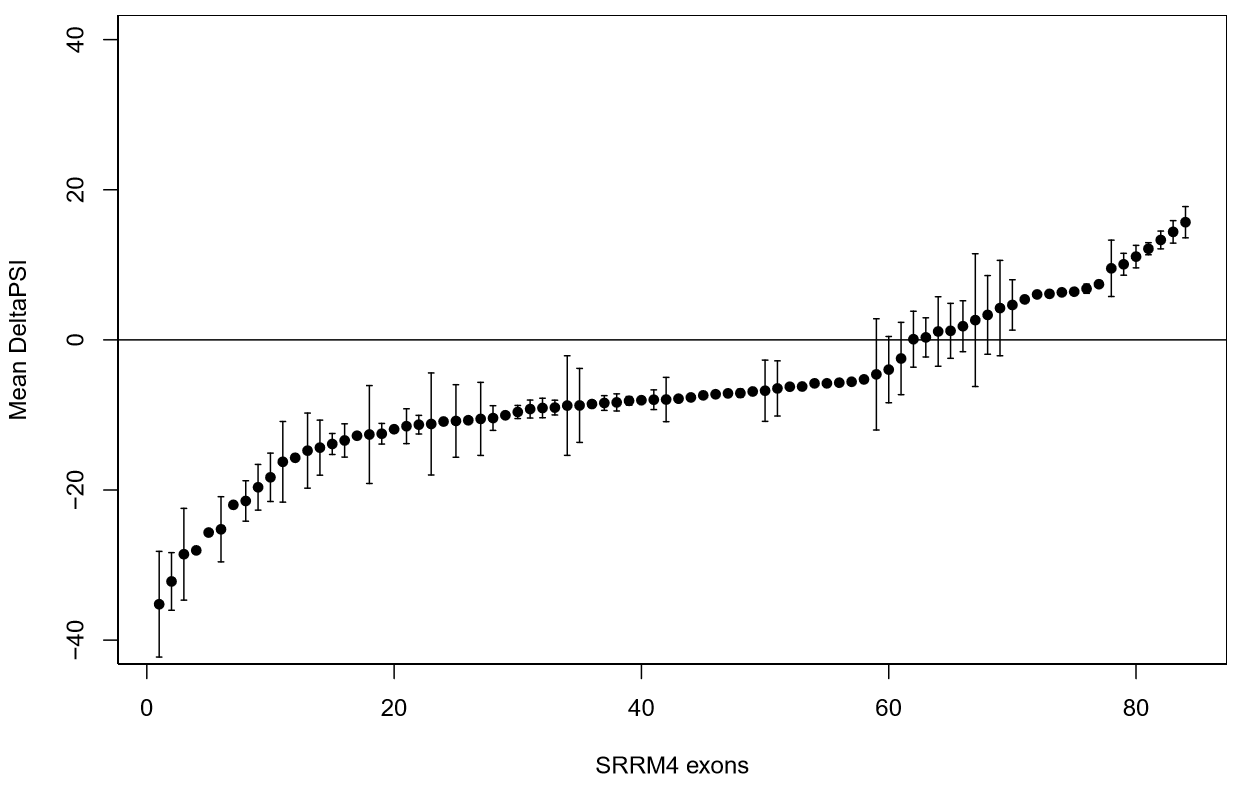

**Fig. S1.** Average ΔPSI (tumor - normal) across tissues for each SRRM4-target exon, demonstrating that more target exons have decreased inclusion in tumors than increased. Only exons with significant changes (q < 0.01, |ΔPSI| ≥ 5) are included.

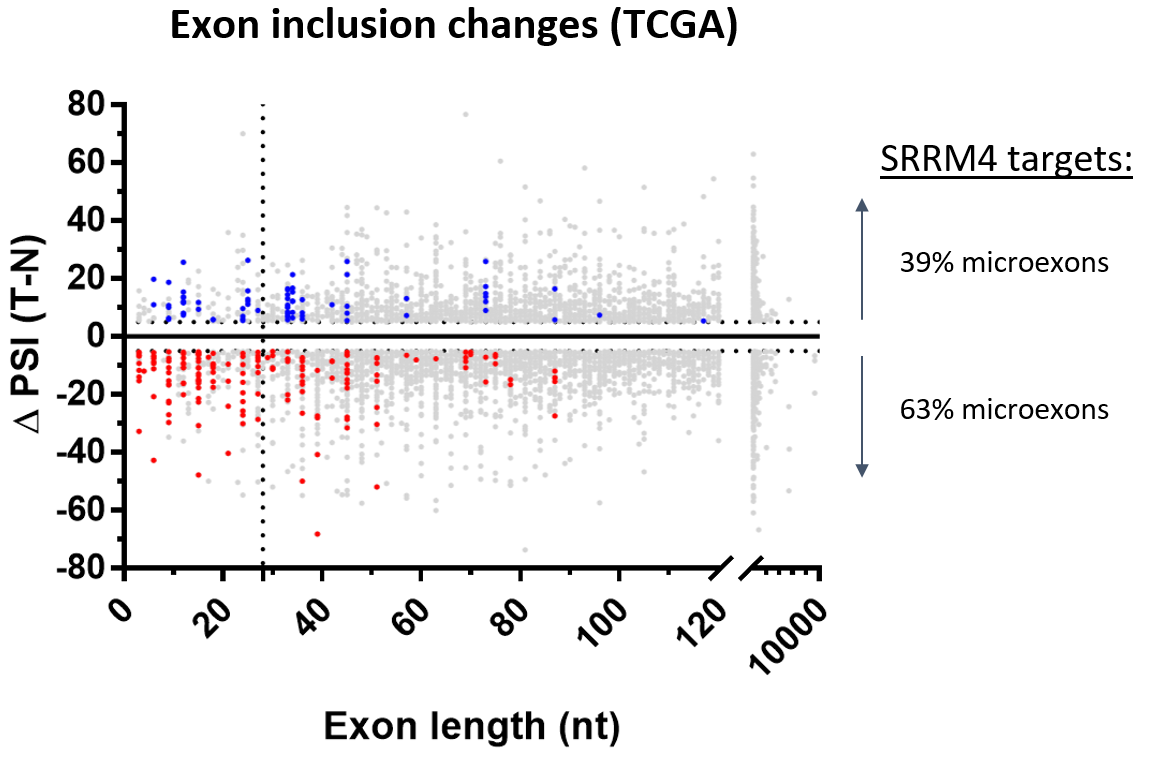

**Fig. S2.** Changes in inclusion level of SRRM4 target exons with respect to exon length. Blue points are SRRM4-target exons increased in tumors (T) vs. normal tissue (N); red are SRRM4-target exons decreased in tumors; grey are non-SRRM4 targets. Only significantly changing exons (q < 0.01) with ΔPSI ≥ 5 are shown. Vertical dotted line separates microexons (≤27 nt) from non-microexons (>27 nt).

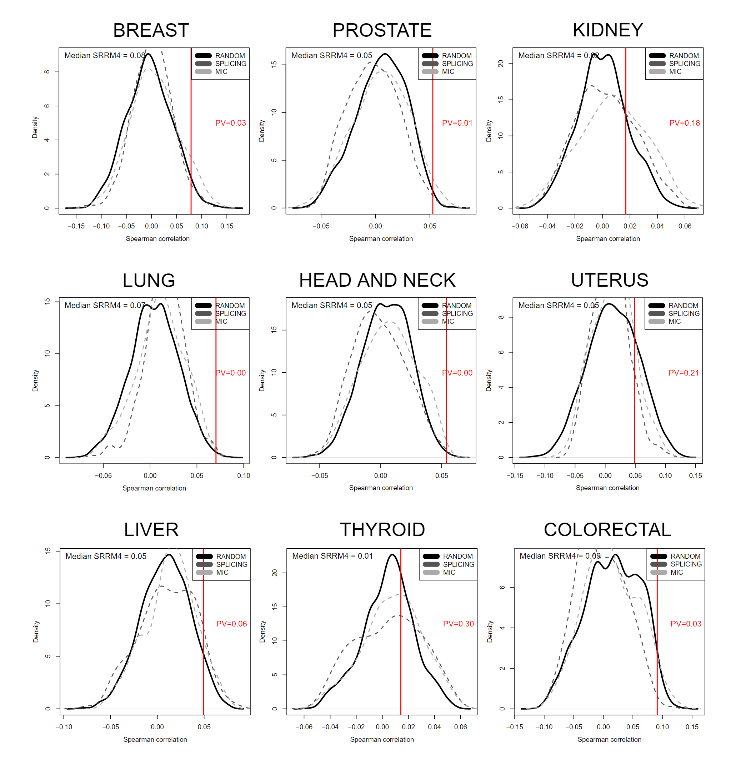
**A	B**

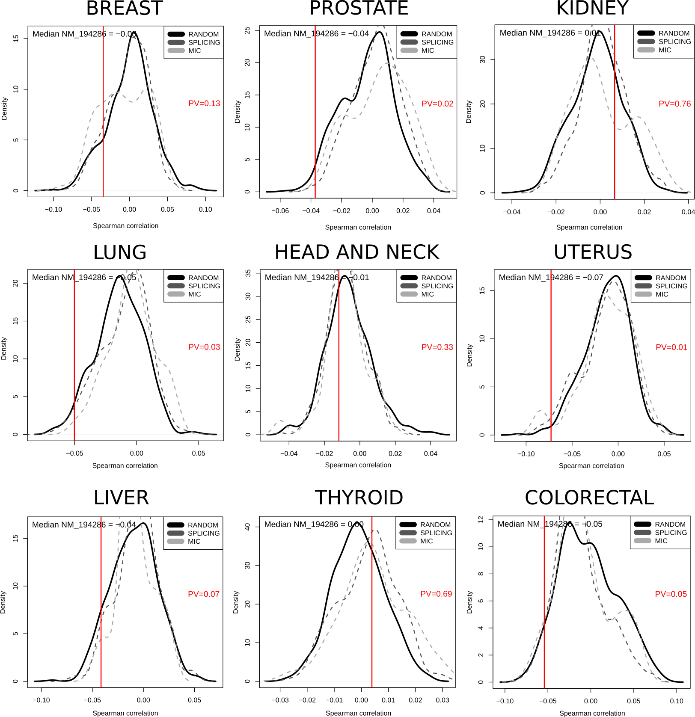

**Fig. S3.** Median Spearman correlation of A) SRRM4 expression and B) SRRM4 methylation with SRRM4-target exons PSI across TCGA tumor samples. The significance of the correlation was determined based on a randomization test within a background of 5000-random-genes background, as well as two other backgrounds of 425 GO-splicing-related genes and 233 microexon-related genes.

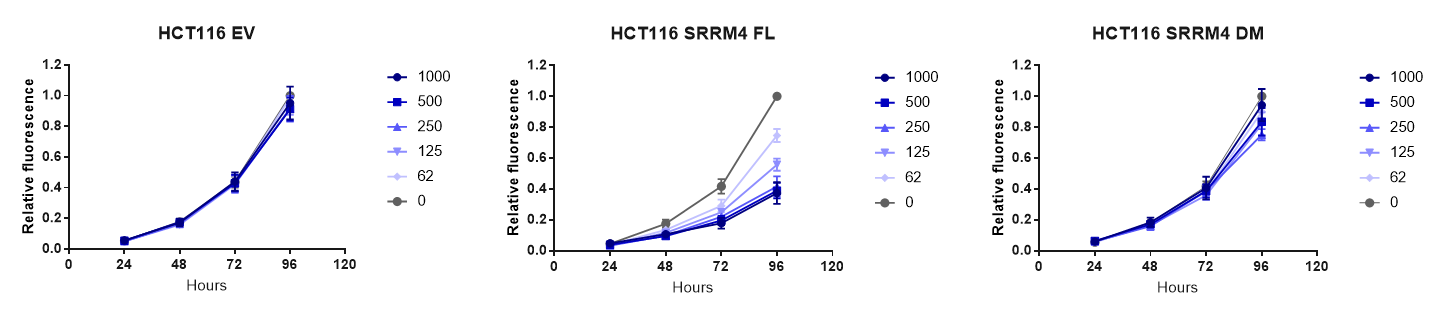

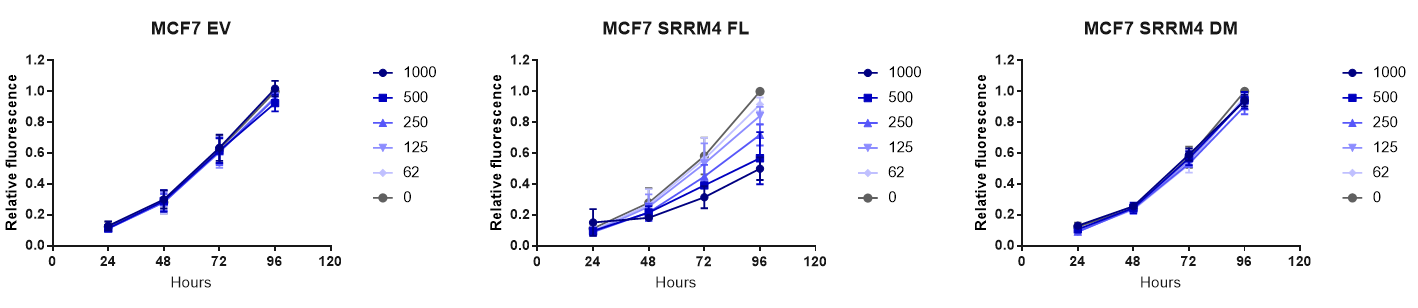

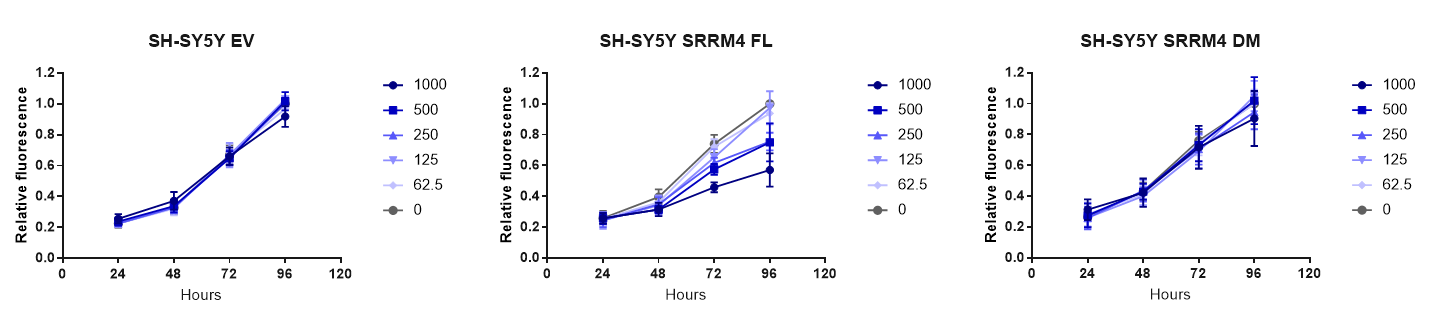

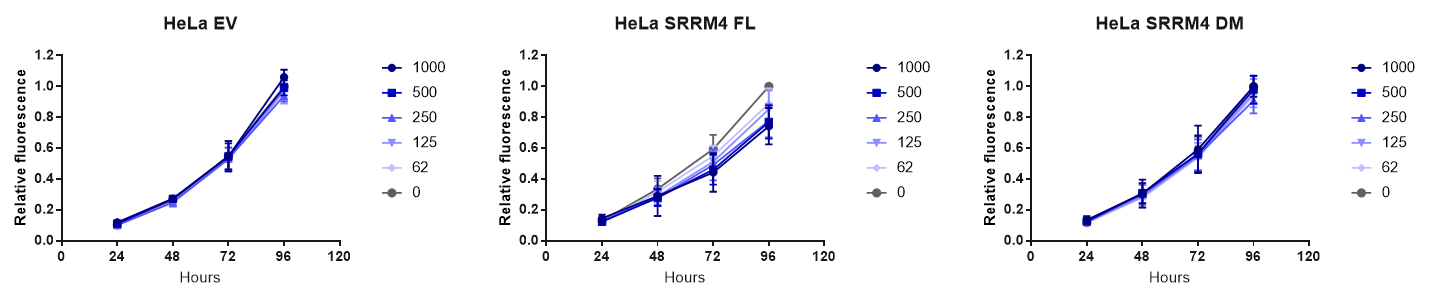

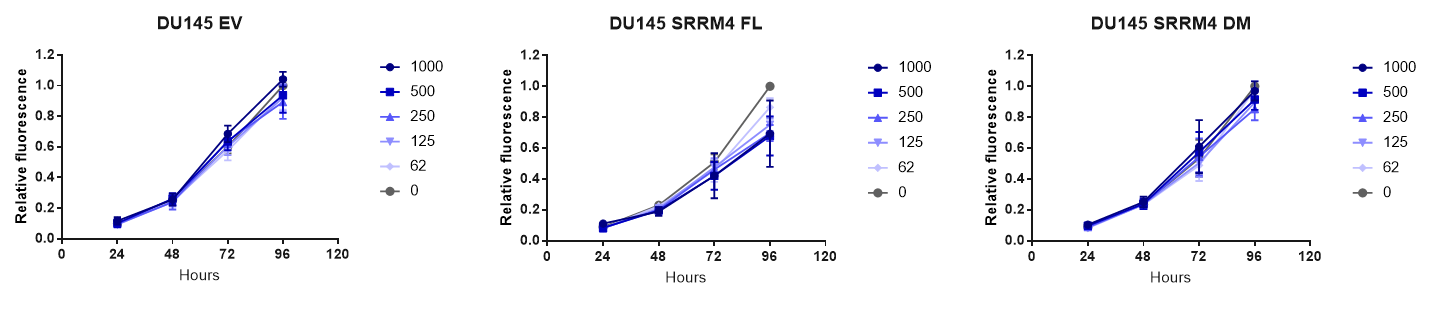

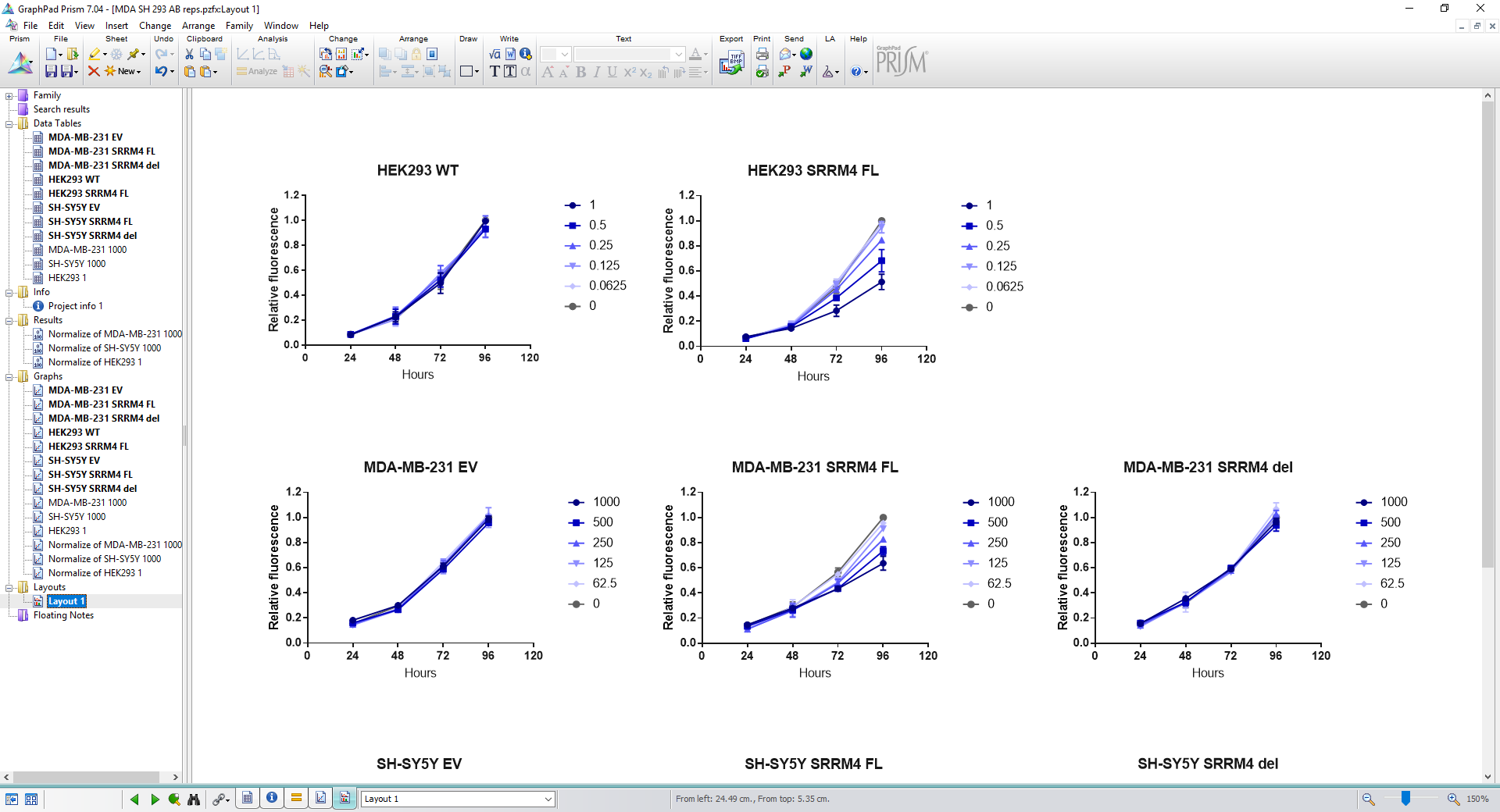

**Fig. S4.** Growth curves of cell lines with inducible SRRM4 expression. Cells were transduced with empty vector (EV), wild-type SRRM4 (WT) or deletion mutant SRRM4 (DM), and treated with indicated concentrations of doxycycline (ng/mL). All lines were generated by lentiviral transduction (see methods) except for HEK293, which was generated by a Flp-In system (Torres-Méndez et al. 2019). Results shown are averages of three independent experiments, and error bars indicate standard error of the mean.

**HCT116**

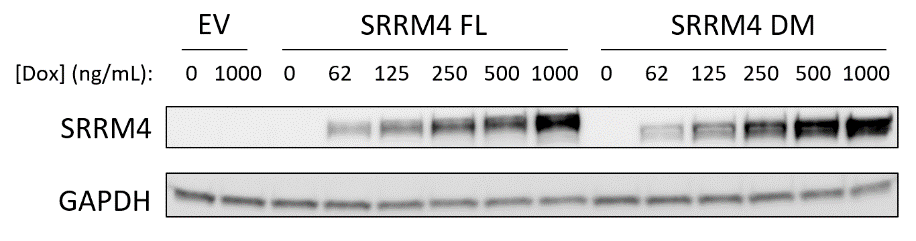

**MCF7**

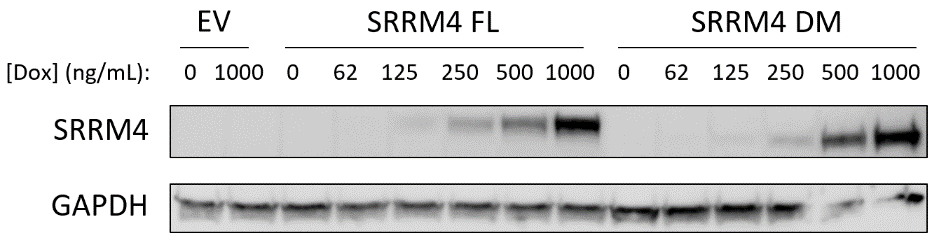

**SH-SY5Y**

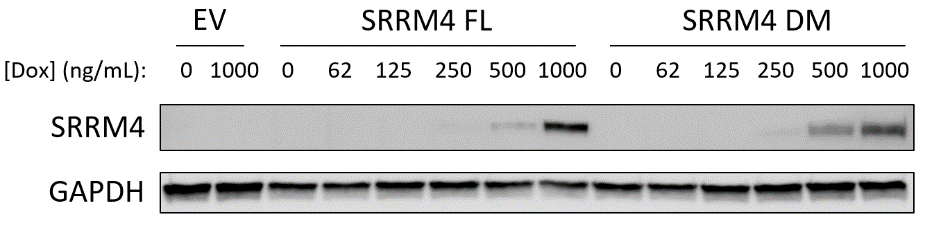

**HeLa**

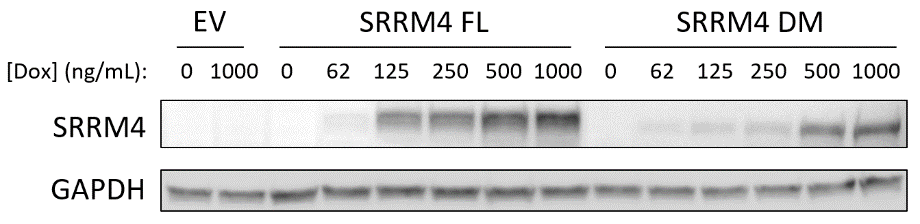

**DU145**

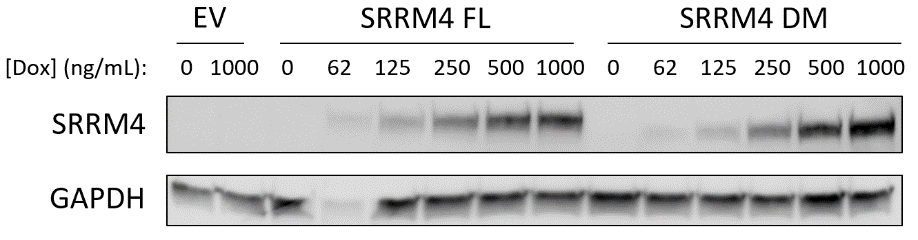

**HEK293**

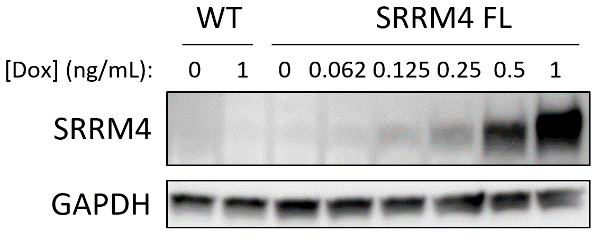

**Fig. S5.** Western blot of SRRM4 (WT) or deletion mutant SRRM4 (DM) induction by doxycycline treatment for 24 hours at the indicated concentrations for the cell lines shown in Fig. S4.

| **TISSUE TYPE** | **TCGA STUDY** | **# TUMOR** | **# NORMAL** |
| --- | --- | --- | --- |
| Breast | BRCA | 1135 | 114 |
| Prostate | PRAD | 505 | 52 |
| Kidney | KIPAN | 901 | 129 |
| Lung | LUAD/LUSC | 1044 | 110 |
| Head and Neck | HNSC | 520 | 44 |
| Uterus | UCEC | 614 | 35 |
| Liver | LIHC | 371 | 50 |
| Thyroid | THCA | 505 | 59 |
| Colorectal | COAD/READ | 669 | 51 |

**Table S1.** Tissues and number of primary tumor and normal solid tissue samples analyzed from TCGA.

**Dataset S1 (separate file).** Differential analysis of the expression of 202 splicing factors using Wilcoxon Ranksum Test.

**Dataset S2 (separate file).** Differential analysis of the methylation of 202 splicing factors using Wilcoxon Ranksum Test.

**Dataset S3 (separate file).** List of significantly differentially included exons between tumor and normal samples from TCGA.

**Dataset S4 (separate file).** List of 314 differentially included exons upon SRRM4 overexpression in HEK293.

**Dataset S5 (separate file).** Inclusion table of SRRM4-target exons in cell lines from the NCI-60.

**Dataset S6 (separate file).** GO-term enrichment analysis of genes with decreased exon inclusion in tumors from TCGA.

**Dataset S7 (separate file).** Gene Expression and Exon Inclusion in 6 cancer cell lines with 24h induction of WT or DM SRRM4.

**Dataset S8 (separate file).** Binomial test sample sizes and p-values.

**Dataset S9 (separate file).** Gene Expression and Exon Inclusion in mouse tumors with or without induction of WT SRRM4.

**Dataset S10 (separate file).** Gene Expression and Exon Inclusion in 2 cancer cell lines with 7 days induction of WT or DM SRRM4.
